# Temporal quantitative proteomics of protein translation and phosphorylation in synaptic plasticity

**DOI:** 10.1101/668434

**Authors:** Charlotte AGH van Gelder, Renske Penning, Lisa Catsburg, Casper C Hoogenraad, Harold D MacGillavry, Maarten Altelaar

## Abstract

At neuronal synapses, activation of metabotropic glutamate receptors (mGluR1/5) triggers a form of long-term depression (mGluR-LTD) that relies on new protein synthesis and the internalization of AMPA-type glutamate receptors. Dysregulation of these processes has been implicated in the development of mental disorders such as autism spectrum disorders and therefore merit a better understanding on a molecular level. Here, to study mGluR-LTD, we integrated quantitative high-resolution phosphoproteomics with the analyses of newly synthesized proteins via bio-orthogonal amino acids (azidohomoalanine) in a pulsed labeling strategy combined with tandem mass tag label-based quantification in cultured hippocampal neurons stimulated with DHPG. We identified several kinases with important roles in DHPG-mGluR-LTD, which we confirmed using small molecule kinase inhibitors. Furthermore, changes in the AMPA receptor endocytosis pathway in both protein synthesis and protein phosphorylation upon LTD were identified, whereby Intersectin-1 was validated as a vital player in this pathway. This study revealed several novel insights into the molecular mechanisms underlying mGluR-LTD and provides a broad view on its molecular basis, which serves as a rich resource for further analyses.

## Introduction

Activation of metabotropic glutamate receptors (mGluRs) initiates a broad array of signaling pathways that collectively modulate the efficiency of neuronal communication. mGluR-dependent signaling has been linked to cognitive functions such as attention, learning and memory, and disrupted mGluR signaling has been implicated in neurological disorders such as Fragile X Syndrome, mental retardation, schizophrenia, addiction and autism spectrum disorders (Bhakar, Dolen et al. 2012, Nicoletti, Bockaert et al. 2011, Niswender, Conn 2010). In particular, group I mGluRs (mGluR1 and mGluR5), generally localized at the postsynaptic membrane, significantly contribute to synaptic function by modulating synaptic excitability, and inducing or facilitating different forms of synaptic plasticity (Bashir, Bortolotto et al. 1993, Bellone, Luscher et al. 2008, Bortolotto, Bashir et al. 1994, Hu, Park et al. 2010, Popkirov, Manahan-Vaughan 2011). Probably the best characterized form of plasticity mediated by group I mGluRs is the long-term depression of synaptic strength referred to as mGluR-LTD (Luscher, Huber 2010). In contrast to NMDA receptor-dependent forms of LTD, the major mechanism of mGluR-LTD expression relies on the rapid and local synthesis of new proteins in dendrites (Huber, Kayser et al. 2000), although not all forms of mGluR-LTD require protein synthesis (Moult, Correa et al. 2008, Nosyreva, Huber 2005). Thus, defining the signaling pathways downstream of mGluR that control translational regulation, and identifying the proteins that are newly synthesized in response to mGluR activation, are important goals in an effort to better understand mGluR-dependent plasticity mechanisms.

Postsynaptic mGluRs canonically link to Gα_q/11_ G-proteins, which activate phospholipase C (PLC) to form diacylglycerol (DAG) and inositol tris-phosphate (IP3). IP3 in turn triggers the release of Ca^2+^ from internal stores, resulting in an increase in the Ca^2+^ concentration and activation of protein kinase C (PKC) (Niswender, Conn 2010). Apart from these pathways, mGluR stimulation has been found to activate a wide range of other downstream effectors, including c-Jun N-terminal kinase JNK1 (Li, X. M., Li et al. 2007), casein kinase 1, cyclin-dependent kinase 5 (CDK5) (Liu, Ma et al. 2001), and components of the ERK-MAPK (Bolshakov, Carboni et al. 2000, Gallagher, Daly et al. 2004, Rush, Wu et al. 2002), and PI3K-Akt-mTOR (Hou, Klann 2004a, Ronesi, Huber 2008, Rong, Ahn et al. 2003) signaling pathways. In particular, induction of these latter two pathways are essential for the expression of mGluR-LTD, mainly because these converge on the regulation of translation initiation factors such as Mnk1, eIF4E and 4EBPs (Banko, Hou et al. 2006, Hou, Klann 2004a).

mGluR-LTD induces an acute wave of new protein synthesis that is required for the long-term reduction in surface α-amino-3-hydroxy-5-methyl-4-isoxazolepropionic acid receptors (AMPAR) underlying the depression of synaptic responses (Huber, Kayser et al. 2000, Waung, Huber 2009). The generation of these “LTD proteins” is directly related to the rate of AMPAR endocytosis, and the rapid synthesis of proteins is required for the internalization of AMPARs and mGluR-LTD (Purgert, Izumi et al. 2014, Nakase, Kobayashi et al. 2015, MacGillavry, Song et al. 2013, Shepherd, Huganir 2007, Davidkova, Carroll 2007, Nadif Kasri, Nakano-Kobayashi et al. 2011, Waung, Pfeiffer et al. 2008, Zhang, Venkitaramani et al. 2008). However, it is unclear what other molecular processes are working in parallel to sustain mGluR-LTD. Thus, even though some of the key mechanisms underlying mGluR-LTD have been identified, characterizing the full repertoire of molecular events that are initiated by mGluR activation would greatly enhance our understanding of mGluR-LTD.

Here, to identify the phosphorylation dynamics initiated by mGluR activation in hippocampal neurons, we applied a phosphoproteomics approach using high-resolution LC-MS/MS. The sensitivity of our approach allowed us to profile multiple time-points over the course of mGluR-LTD. In addition, to identify newly synthesized proteins in response to mGluR activation, we used an azidohomoalanine (AHA) labeling strategy (Eichelbaum, Winter et al. 2012) in combination with tandem mass tag (TMT) labeling for accurate quantification of translated proteins. Based on the observed phosphorylation dynamics we identified several kinases important in the regulation of DHPG-mGluR-LTD, which we confirmed using specific kinase inhibitors. Furthermore, we uncovered a broad spectrum of protein synthesis and phosphorylation dynamics in the AMPA receptor endocytosis pathway upon LTD. We thereby highlight several novel insights into the mechanism of mGluR-LTD, and validated Intersectin-1 (Itsn1) to play an important role in AMPAR trafficking during mGluR-LTD.

## Results

### Protein synthesis upon activation of mGluR-LTD via DHPG

To induce mGluR-LTD in primary hippocampal cultures, neurons were stimulated with DHPG, a specific agonist of group I mGluRs. We confirmed that this induced a reduction in surface GluA1 expression (Figure S1A) as has been established before (Anggono, Huganir 2012, Luscher, Huber 2010). To identify proteins that are synthesized *de novo* in response to mGluR activation by DHPG, we used a pulsed-AHA approach (Dieterich, Link et al. 2006, Eichelbaum, Winter et al. 2012), where cultured hippocampal neurons were stimulated with DHPG in the presence of the bio-orthogonal methionine analogue AHA to label newly synthesized proteins. To profile the temporal induction of protein synthesis, neurons were harvested and lysed at different time points up to 90 minutes after a 5-minute DHPG stimulation (Figure 1A). AHA-incorporated proteins were consecutively enriched via click-chemistry, digested by a combination of Lys-C and trypsin and analyzed using high-resolution nanoLC-MS/MS. For each studied time point, control experiments were included where either AHA was supplemented to unstimulated neurons, to define the set of proteins that are truly being translated in response to DHPG, or neurons were stimulated with DHPG in methionine-supplemented media in the absence of AHA, to control for non-specific binding during the enrichment process.

**Figure 1.**
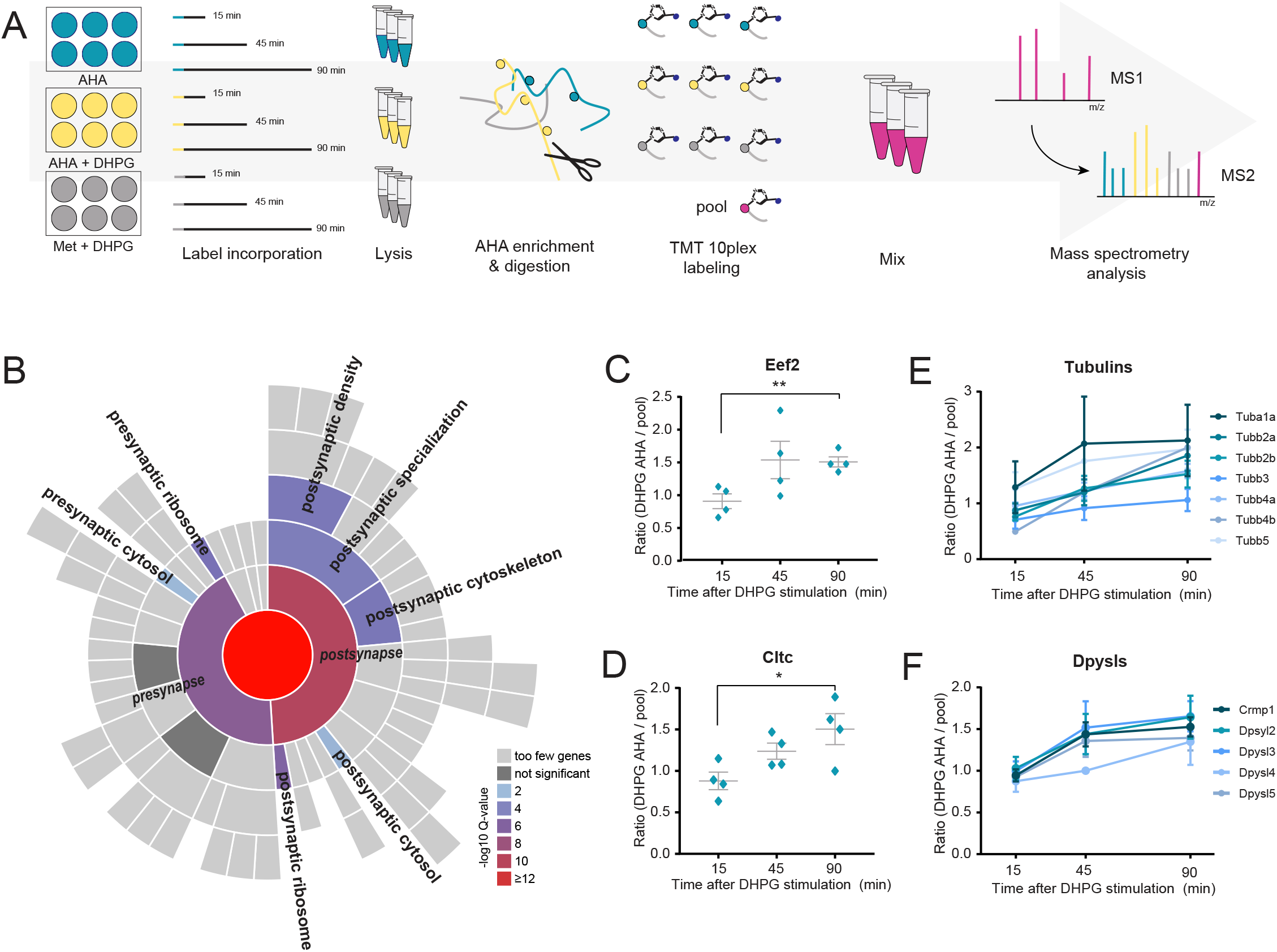
Identification of protein translation following mGluR-LTD induction. (A) Workflow: neurons were stimulated with DHPG for 5 minutes in combination with methionine (negative control) or the methionine substitute AHA. An extra control was included in which neurons were not stimulated with DHPG but were supplemented with AHA. After DHPG removal, translation was followed for a total of 15, 45, and 90 minutes. Newly synthesized proteins were enriched, digested and labeled with TMT10plex. (B) SynGO cellular location enrichment analysis of newly translated proteins (1% FDR) revealed enrichment of postsynaptic over presynaptic localization. (C) Induction of mGluR-LTD using DHPG leads to identification of proteins involved in translation, such as Eef2, as well as (D) in receptor endocytosis, such as clathrin, showing increasing protein abundance over time. (E) This enrichment and labeling approach reducibly identifies multiple members of the same protein families during mGluR-LTD, including subunits of both alpha and beta tubulins and (F) Dpysls. Data are represented as mean±SEM, * p<0.05, p<0.01.

For further analysis, only proteins that were identified in at least two biological replicates of at least one experimental condition were considered (273 proteins). Furthermore, only DHPG specific and AHA enriched proteins that had a higher relative expression than in the control conditions in at least one of the time points were considered (266 proteins, Table S1). Notably, less than 3% of the identified proteins did not fulfill the methionine control criterion, and less than 10% of all newly translated proteins were also identified in the AHA control condition without DHPG stimulation and were considered to be background. Also, it was assuring that the number of newly synthesized proteins increased over time, and the overlap between these proteins was high between all experimental time points (Figure S2), highlighting the sensitivity of the experimental approach. Indeed, GO enrichment analysis against a full brain background using SynGO (Koopmans, van Nierop et al. 2019) confirmed enrichment of postsynaptic over presynaptic localization, as well as a clear enrichment in, among others, postsynaptic cytoskeleton and ribosome (Figure 1B). Among the newly synthesized proteins are hallmark mGluR-LTD proteins involved in translation, such as translation initiation factor 4a (Eif4a2) and elongation factor 2 (Eef2) (Waung, Huber 2009, Gladding, Fitzjohn et al. 2009), which demonstrates the effectiveness of this experimental set up to identify protein translation upon mGluR-LTD induction (Figure 1C).

The induction of mGluR-LTD ultimately results in augmented internalization of AMPARs (Davidkova, Carroll 2007, Huber, Kayser et al. 2000, Luscher, Huber 2010, Waung, Pfeiffer et al. 2008, Waung, Huber 2009, Zhang, Venkitaramani et al. 2008). Accordingly, we found many newly synthesized proteins with functions related to receptor endocytosis and recycling. Examples include the adaptor protein complex AP-2 subunit (Ap2a1), the GTPase Cdc42, and the neuronal migration protein doublecortin (Dcx), which is a known interactor of clathrin adaptor complexes (Friocourt, Chafey et al. 2001) (Table S1). In line with this observation, we identified DHPG-dependent translation of the heavy chain of clathrin (Cltc), a key protein for the formation of coated vesicles, which was steadily being translated over all three time points (Figure 1D).

In line with the findings that reorganization of the actin cytoskeleton underlies the expression of mGluR-LTD (Sanderson, Hogg et al. 2016), and is for instance thought to cause the changes in spine morphology that are associated with mGluR-LTD (Zhou et al., 2011), we observed several proteins with known roles in synaptic plasticity processes, including stathmins (Stmn1 and Stmn2), profilin (Pfn2), and neuromodulin (Gap43), as well as cofilin-2 (Cfl2) (Table S1). Interestingly, apart from these actin regulators we also observed profound translation of microtubule-associated proteins (Map2, Map6, Mapre1, Map1lc3b, and Mapt), as well as numerous subunits of both alpha and beta tubulins over all time points (Figure 1E).

Interestingly, we also identified and measured the translation of several protein families and complexes over time, such as several proteasomal subunits, ribosomal proteins, 14-3-3 proteins, tubulins, and dihydropyrimidinase-related proteins, which are also known as the collapsing response mediator protein family (CRMPs) (Figure 1F). These versatile proteins are involved in a variety of developmental and plasticity-related brain processes, and have been shown to also localize in the PSD (Bayés, Collins et al. 2012). Interestingly, the CRMP family of proteins were recently identified as some of the most stable, long-lived proteins in neurons (Heo, Diering et al. 2018).

To generate a more in-depth overview of the type of proteins being translated, and which processes they represent, we performed a k-means interaction-based clustering analysis using the protein-interaction database STRING (Table S1) (Szklarczyk, Franceschini et al. 2014). GO analysis of these resulted in the enrichment of six biological processes, visualized using the geneMANIA app in Cytoscape (Montojo, Zuberi et al. 2010). This interaction-based clustering approach highlights interactions between proteins as identified by, among others, physical interactions, co-expression data and co-localization data. With these combined data, we can identify groups of proteins with similar functions in relevant biological processes underlying the expression of mGluR-LTD. Figure 2 displays the interaction network with representative proteins of each of these processes, as well as their z-score normalized expression over all pooled experimental conditions. As expected, we identify several protein clusters with an enrichment in biological processes related to protein translation. The first cluster contains proteins involved in the processing of mRNA, and includes several splicing factors (Srsf2 and Srsf3) and ribonucleoproteins (*e.g.* Snrnp70), as well as binding proteins (Nono and Pcbp2). The second cluster, with a clear enrichment in proteins regulating translation, includes translation initiation factor Eif4a2, and elongation factors (Eef1g, Eef2, and Tufm), as well as multiple ribosomal proteins, both from the 40S (*e.g.* Rps2 and Rps7) and 60S ribosome (*e.g.* Rpl3 and Rpl22). This suggests that mGluR-LTD induction initiates the formation of new ribosomes, presumably to meet the demands of an increased rate of protein synthesis. Cluster 3 contains proteins involved in protein folding, including several heat shock proteins (*e.g.* Hsp90aa1 and Hsp90ab1), as well as several subunits of the t-complex (*e.g.* Cct2 and Cct5), which are molecular chaperones in protein folding (Frydman 2001). Recently, a dynamic interplay between protein translation and degradation has been described to be crucial for mGluR-LTD (Klein, Castillo et al. 2015, Ramachandran, Margolis 2017). Inhibition of the proteasome rescued impairments in mGluR-LTD induction caused by blockage of protein translation (Klein, Castillo et al. 2015), suggesting that a fine coordination between protein translation and degradation of proteins by the UPS is key for the efficient induction of mGluR-LTD. In line with this, we observed translation of many proteins involved in protein degradation in cluster 4, including several subunits of the proteasome (*e.g.* Psma2, Psma3, Psmb1 and Psmb7). Finally, we identified two protein clusters with a similar enrichment in GO terms, containing proteins involved in the organization of cellular components (cluster 5) and, more specifically, cytoskeleton organization (cluster 6). While both clusters contain multiple tubulins, cluster 5 also contains several 14-3-3 proteins, which have been studied extensively in relation to their function in the regulation of actin filaments (Sluchanko, Gusev 2010). In addition to tubulins, cluster 6 also contains several actin regulating proteins (*e.g.* Arpc1a and Wdr1).

**Figure 2.**
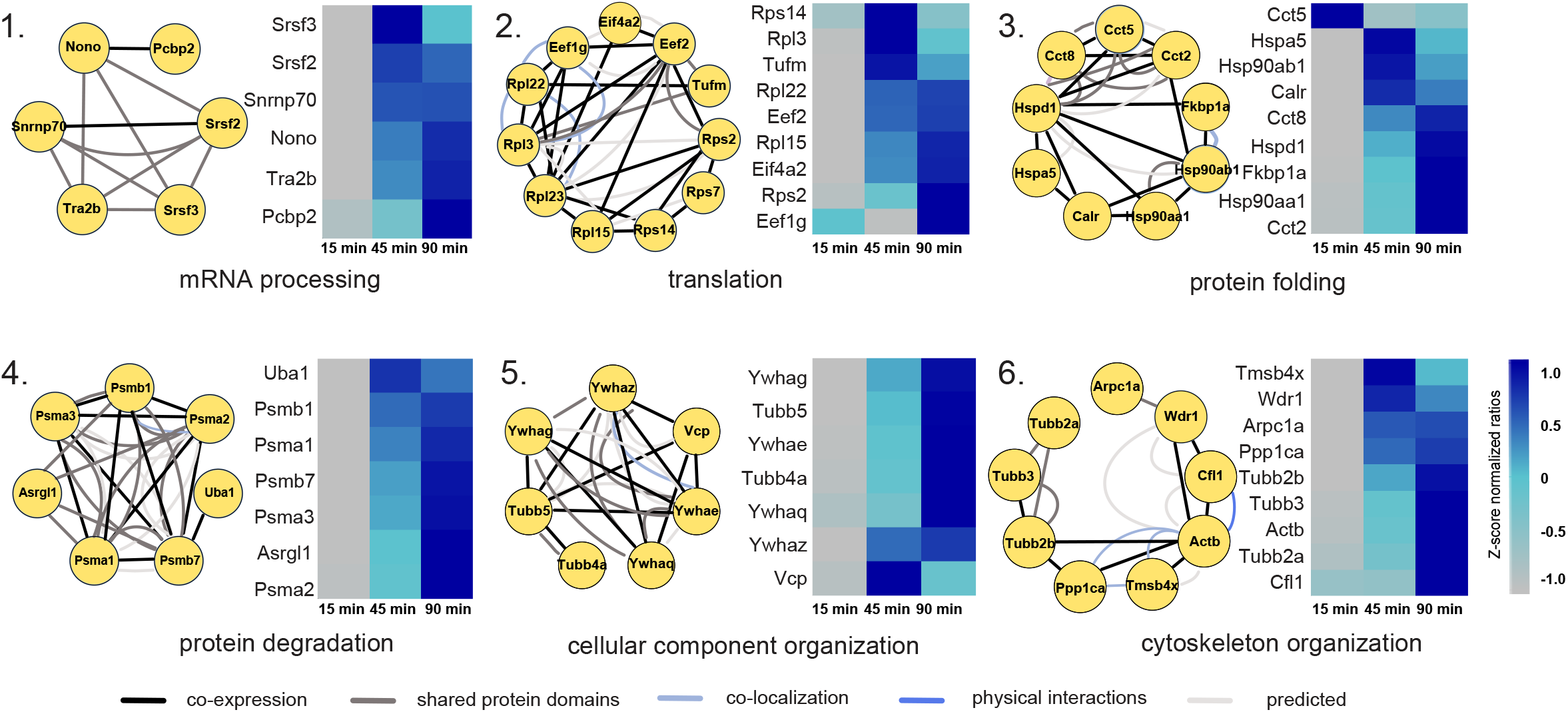
Interaction-based protein clusters with distinct biological processes involved in mGluR-LTD. GO enrichment analysis of k-means based interaction clusters resulted in six main clusters with a clear enrichment in a biological process. Proteins from each cluster are displayed with their known interaction profiles. Heatmaps represent the z-score normalized ratio DHPG AHA / pool over the three measured time points.

### Phosphoproteomics of mGluR-LTD activation in primary neurons reveals defined clusters of phosphosite regulation with a strong synaptic signature

Due to the low stoichiometry of phosphorylation events in the proteome, phosphoproteomics analysis involves dedicated enrichment strategies. These strategies typically require milligrams of protein input material (Humphrey, Azimifar et al. 2015, Villen, Gygi 2008, Zarei, Sprenger et al. 2011), hampering analysis of phosphorylation dynamics in primary neurons. Recently, we have shown that automated phosphopeptide enrichment using Fe(III)-IMAC cartridges on a Bravo AssayMap platform allows sensitive and reproducible enrichment of several thousands of unique phosphopeptides starting with only 1-10 µg of protein input material. Moreover, we showed that from a single neuronal culture plate, 200,000 cells delivering ∼50 µg of protein, we could identify biological relevant phosphorylation events among the ∼7,000 observed phosphosites (Post, Penning et al. 2016). This now allowed us to study phosphorylation events on multiple time-points during mGluR-LTD in primary hippocampal neurons, without the need for combining extensive amounts of input material. The proteomics analysis of Fe(III)-IMAC enriched phosphorylated peptides was performed after stimulating rat hippocampal neurons for 0, 5, 10 or 20 minutes with DHPG. The applied workflow is outlined in Figure 3A. This phosphoproteomics screen resulted in the identification of 17,556 phosphosites with a localization probability >0.75, of which 5,423 could be quantified in at least two biological replicates in at least one experimental condition and were used for subsequent analysis (Table S2). Phosphopeptide abundance showed a normal distribution (Figure S3A) and a high degree of overlap between the identified phosphoproteins from both the 5, 10 and 20 minutes DHPG-stimulated neurons could be observed compared to control (Figure S3B). The distribution of phosphosites on serine, threonine and tyrosine residues is in line with previous publications (Humphrey, Azimifar et al. 2015, Post, Penning et al. 2016, Ye, Zhang et al. 2010) (Figure S3C). The reproducibility of the experimental procedure was assessed between biological replicates, and between the different time points. A high degree of correlation between the biological replicates could be observed, suggesting that a small, but distinct subset of signaling pathways is activated by DHPG, consistent with the notion that DHPG activates local dendritic signaling pathways, rather than a robust, cell-wide response. Nevertheless, the DHPG treated samples could still be clearly distinguished from the control samples (Figure S3D). Furthermore, SynGO enrichment analysis of biological function compared to a full brain background resulted in a significant enrichment for PSD organization, regulation of postsynaptic neurotransmitter receptor activity and localization, and organization of the postsynaptic actin skeleton (Figure S3E).

**Figure 3.**
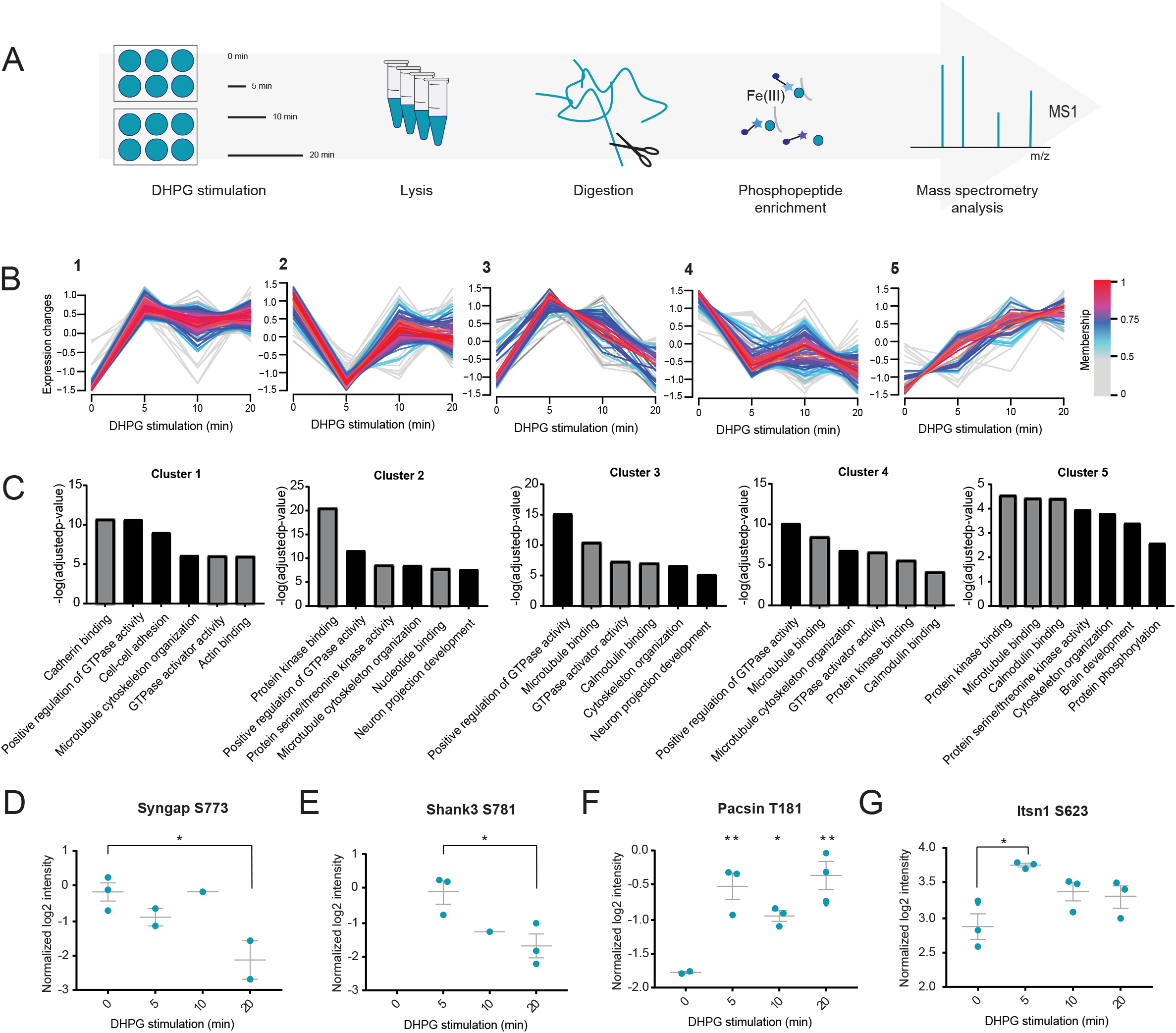
Quantitative phosphoproteomics of mGluR-LTD in hippocampal neurons stimulated with DHPG. (A) Quantitative phosphoproteomics workflow: samples were taken at 0, 5, 10 and 20 minutes after the addition of DHPG. (B) Unsupervised clustering reveals five distinct clusters for the regulated phosphosites. (C) GO-term enrichment analysis for molecular function (grey) and biological process (black). (D-G) Normalized log2 intensities of regulated phosphosites upon DHPG stimulation. Data are represented as mean±SEM, * p<0.05, ** p<0.01.

To identify phosphosites regulated over time, a multiple-samples ANOVA was performed, resulting in a total of 504 phosphosites significantly regulated in response to DHPG (*p*<0.05). Unsupervised fuzzy clustering of these regulated phosphosites revealed five distinct clusters (Figure 3B) (Rigbolt, Vanselow et al. 2011). The phosphosites grouped in cluster 1 are characterized by an increasing trend in phosphorylation and are enriched in multiple GO terms compared to the full phosphoproteome, such as actin binding and microtubule cytoskeleton organization (Figure 3C). In cluster 2, showing a distinct pattern starting with immediate de-phosphorylation upon mGluR-LTD induction followed by re-phosphorylation at later time points, we observe enrichment in processes involved in signaling, such as protein kinase binding and serine/threonine kinase activity. Cluster 3, containing proteins involved in cytoskeleton organization and microtubule binding (*e.g.* Map1b and Mapt), shows initial phosphorylation followed by dephosphorylation at the later time points. Both clusters 4 and 5 show a steady trend in phosphorylation over all time points, where proteins are either steadily dephosphorylated over time (cluster 4), or phosphorylated over time (cluster 5). Both clusters are enriched for proteins associated with microtubule binding (*e.g.* Map2, Map4 and Macf1), small GTPase regulator activity, and more generally protein phosphorylation.

### Phosphorylation dynamics in relation to DHPG-activated mGluR-LTD

Regulation of major signaling pathways by kinases and phosphatases underlies various dynamic processes in cellular functioning, including mGluR-LTD (Collingridge, Peineau et al. 2010, Gallagher, Daly et al. 2004, Gladding, Fitzjohn et al. 2009, Hou, Klann 2004a, Moult, Gladding et al. 2006). A selection of several kinases can influence important nodes of these signaling pathways. One kinase shown before to be important in mGluR-LTD is Ca^2+^/calmodulin-dependent protein kinase II (CaMKII) (Bernard, Castano et al. 2014, Mockett, Guevremont et al. 2011). We found subunit CaMKIIβ to be dephosphorylated in our dataset at S315 following DHPG stimulation. Interestingly, this phosphorylation site is located in its F-actin binding domain and is potentially regulated by autophosphorylation (Kim, Lakhanpal et al. 2015). Next to CaMKII, we observed dephosphorylation, and thus activation, of Eef2 at T57, the major target of the Ca^2+^/calmodulin-dependent protein kinase eEF2K (Hizli, Chi et al. 2013). Interestingly, one hour after DHPG stimulation, Eef2 phosphorylation has been shown to result in mGluR-LTD related protein translation of Arc/Arg3.1 and inhibition of global protein translation (Park, Park et al. 2008).

Next to protein translation, several proteins involved in protein degradation showed regulation by phosphorylation upon DHPG stimulation. Among these are the T273 phosphosite of the Psmd1, and T9 of the Psmd2 regulatory subunits of the 26S proteasome, several other proteasome subunits (Psma5 and Psmc3), and several ubiquitin-related enzymes.

Multiple integral proteins of the PSD showed regulation at the phosphorylation level. For instance, the protein SynGAP1, an important negative regulator of AMPAR insertion at the membrane of the PSD (Rumbaugh, Adams et al. 2006), was significantly dephosphorylated over time at S773 (Figure 3D). Phosphorylation of the S773 site alone has been shown to inhibit GAP activity, while concurrent phosphorylation of both S773 and S802 increased GAP activity (Walkup, Washburn et al. 2015). GAP activity is necessary for the inactivation of Ras and Rap, which are involved in AMPAR trafficking (Walkup, Washburn et al. 2015). Our data thus suggest that dephosphorylation of SynGAP1 at S773 in response to DHPG changes the Ras/Rap activation balance, perhaps promoting AMPAR endocytosis.

Significant changes in phosphorylation status were also found in several PDZ domain-containing proteins (Table S2). We observed alterations in phosphorylation in prominent synaptic scaffolding proteins such as Dlg2 (PSD-93), Dlg3 (SAP-102), Shank2 and Shank3. The latter was found to be phosphorylated already five minutes after mGluR-LTD induction at S781 (Figure 3E), a site that has not previously been identified in rat, but was recently shown to be induced by an LTP protocol in mouse PSD fractions (Li, J., Wilkinson et al. 2016). Other regulated PDZ proteins include microtubule-associated serine/threonine-protein kinase 2 (Mast2) at two distinct phosphorylation sites, as well as Rho GTPase-activating proteins Arhgap21 and Arhgap23. Also identified to be phosphorylated, but not found to be significantly regulated during mGluR-LTD, is the PDZ scaffold protein GRIP, which is involved in AMPAR trafficking (Derkach, Oh et al. 2007, Henley, Wilkinson 2016).

Multiple proteins involved in endocytosis show significant regulation at the phosphorylation level, including Syndapin-1 (Pacsin1) and β-Pix (Arhgef7). Syndapin-1 becomes phosphorylated at the T181 site after mGluR-LTD induction (Figure 3F). This phosphosite of Syndapin-1 is located in the F-Bar domain of the protein and is important in neuronal membrane tubulation. The F-bar domain is involved in lipid binding and cytoskeleton reorganization (Quan, Xue et al. 2012). However, this specific phosphosite was not shown to be involved in the regulation of activity-dependent bulk endocytosis. More recently, it was shown that Syndapin-1 also interacts directly with Pick1 via its F-bar domain and that this interaction is important for AMPAR endocytosis in NMDAR-related cerebellar LTD (Anggono, Koc-Schmitz et al. 2013). This might suggest a potential role for this T181 phosphosite in the Syndapin-1 and Pick1 binding during mGluR-LTD, potentially influencing AMPAR endocytosis. β-Pix is a guanine nucleotide exchange factor and binds the p21–activated kinase Pak1. The site S71 of β-Pix is strongly conserved and has a possible role in its guanine exchange function (Mayhew, Jeffery et al. 2007). In our experiment, the phosphorylation pattern of S71 belongs to cluster 2, where after an initial rapid dephosphorylation, the phosphorylation state after 20 minutes of DHPG stimulation returns close to its initial level. Furthermore, two phosphosites (S623 and S1134) of Itsn1, a protein linked to receptor internalization, were significantly regulated upon DHPG stimulation (Figure 3G). In neurons, Itsn1 was suggested to be involved in presynaptic vesicle recycling (Pechstein, Bacetic et al. 2010, Yu, Chu et al. 2008), but was also shown to have a postsynaptic function in AMPAR internalization in invertebrates (Glodowski, Chen et al. 2007). The exact role of these phosphosites is still unknown, also in mGluR-LTD, which makes Itsn1 an interesting target for further study.

### Activated kinases upon mGluR-LTD

To assess if certain kinases are specifically involved in the phosphorylation events underlying mGluR-LTD, we next performed a phosphorylation site consensus motif analysis of the regulated phosphosites. This allowed us to extract multiple phosphorylation motifs (Figure S4) resulting in four distinct typical kinase motif sequences linked to regulated proteins in our dataset (Figure 4A). The first and most pronounced motif is the proline directed motif at the +1 position, which is a known recognition motif of the cyclin-dependent kinases (CDKs). Moreover, we identified the threonine directed TPxK motif, which is a known substrate binding site of Cdk5 (Sharma, Steinbach et al. 1999). The third motif is the RxxS motif, which is part of the known targeting sequence for the MAPK-activated protein Kinases (MKs) (Gaestel 2006, Stokoe, Caudwell et al. 1993). A double MK2/3 knockout mouse model showed impaired mGluR-LTD and GluA1 endocytosis, indicating a regulatory role for these kinases in the process. This RxxS motif is also described as a consensus motif for CaMKII kinases, as is the KxxS motif, which is also visible as the fourth most dominant motif. To experimentally validate the identification of these motifs we used well-characterized pharmacological inhibitors to specifically block the activity of CDKs (roscovitine), or CaMKII (KN-93) before the induction of mGluR-LTD with DHPG in hippocampal cultures. Confirming the predictions, we found that pre-incubation with roscovitine or KN-93 blocked the DHPG-induced reduction in surface GluA1 levels (Figure 4B,C). On the other hand, pre-incubation with staurosporine, blocking PKC activity did not prevent mGluR-induced GluA1 internalization. These experiments thus confirmed our prediction that the activity of CDKs and CaMKII underlie the expression of mGluR-LTD.

**Figure 4.**
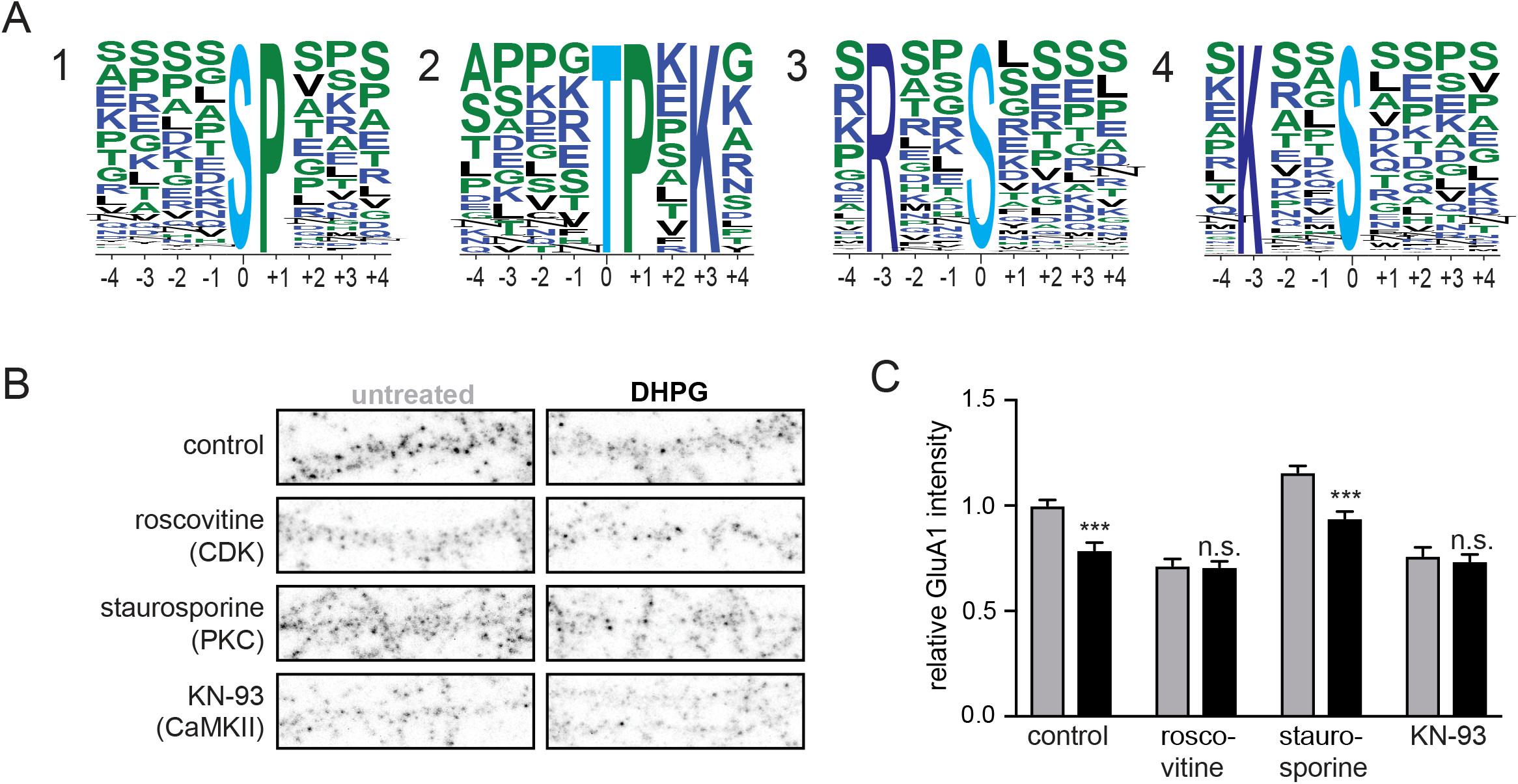
Kinases involved in DHPG-activated mGluR-LTD. (A) MotifX sequence motifs of regulated phosphosites (*p*<0.01) implicated in mGluR-LTD. (B). Immunostaining of GluA1 subunits at the cell surface after control or DHPG treatment, pre-incubateed with control (untreated, N=19)) or 3 different kinase inhibitors (Roscovitine (CDKs, N=15), staurosporine (PKC, N=12) and KN-93 (CaMKII, N=12)). (C) Relative quantification of cell surface GluA1 intensity; pre-incubation with roscovitine (N=13) or KN-93 (N=12) blocked DHPG-induced reduction in surface GluA1 levels in contrary to staurosporine (N=12) pre-incubation which did not prevent mGluR-induced GluA1 internalization. Data are represented as mean±SEM. *** p<0.001, n.s. not significant.

### Newly translated proteins regulated by phosphorylation

Fifteen proteins were found to be both newly synthesized and regulated by phosphorylation upon DHPG stimulation (Table S2). These proteins include several proteins involved in translation and protein synthesis and degradation, as well as microtubule-associated protein 6 (Map6), neuronal migration protein doublecortin (Dcx) and previously discussed elongation factor (Eef2). It also contains neuromodulin (Gap43), a PKC substrate that is known for its presynaptic role in NMDAR-mediated LTD (Ramakers, G. M., Heinen et al. 2000). More recent studies however, have shown that neuromodulin is also abundantly expressed postsynaptically, where it has to be cleaved by Caspase-3 to regulate LTD (Han, Jiao et al. 2013). Interestingly, two of its regulated phosphosites upon mGluR-LTD induction (T95 and S96) are located in close proximity of a Caspase-3 cleavage site. Dephosphorylation of two amino acids in this cleavage domain could very well result in a conformational change, making it more accessible to caspase cleavage. Integration of the phosphoproteomics and protein synthesis datasets, in combination with the data generated from the kinase motif analysis, resulted in a global view of molecular events underlying mGluR-LTD (Figure 5).

**Figure 5.**
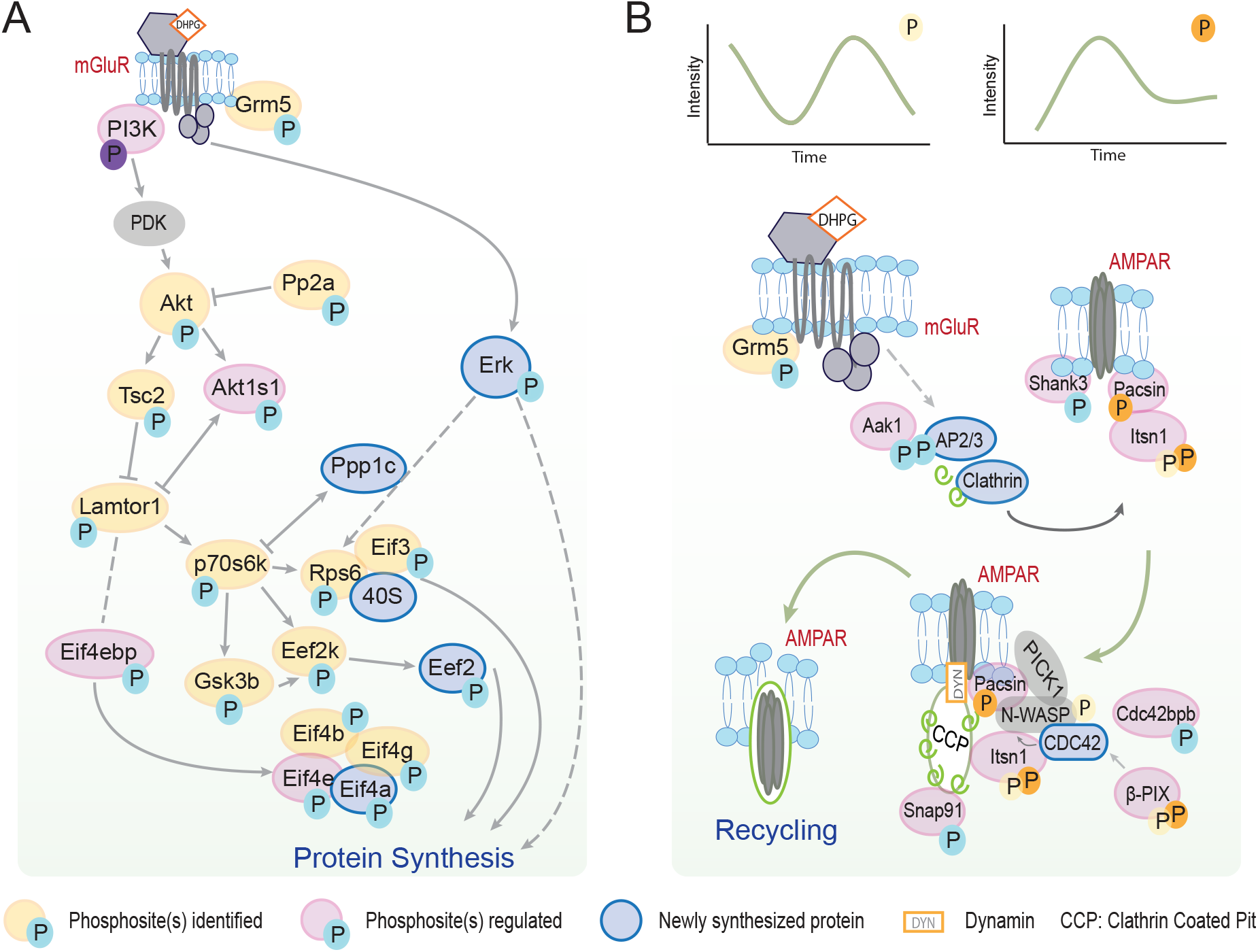
Pathways involved in DHPG-induced mGluR-LTD. (A) Detailed overview of the molecular pathways involved in DHPG-induced mGluR-LTD identified in the (phospho)proteomics experiments, initializing protein translation necessary for mGluR-LTD maintenance. (B). Visualization of phosphorylated proteins shows the activity of several signaling cascades facilitating and activating AMPAR recycling and internalization. The relative quantification of these phosphorylation events (upper panel) shows distinct phosphorylation patterns over time.

### Itsn1 is essential for DHPG-induced AMPAR internalization

In the phosphorylation data we identified Itsn1, a guanine exchange factor (GEF) for the GTPase Cdc42 (Hussain, Jenna et al. 2001), to be regulated at two phosphosites (S623 and S1134) upon DHPG stimulation (Figure 3D). To test whether Itsn1 also has a functional role in mGluR-LTD, we transfected neurons with a miRNA-based knockdown construct targeting both the long and short forms of Itsn1 (mirItsn1) (Thomas, S., Ritter, Verbich, Sanson, Bourbonniere, McKinney, and McPherson 2009a). Immunostaining of endogenous Itsn1 in control neurons showed a punctate pattern, as described before (Thomas, S., Ritter, Verbich, Sanson, Bourbonniere, McKinney, and McPherson 2009a), and confirmed significant depletion of Itsn1 in mirItsn1-transfected neurons (Figure 6A,B). Interestingly, surface GluA1 expression was significantly reduced in Itsn1 knockdown neurons under basal conditions, indicating that Itsn1 is involved in the regulation of AMPAR surface expression. Furthermore, we found that the reduction in GluA1 surface levels in response to DHPG was severely affected in Itsn1 knockdown neurons (Figure 6C), indicating that Itsn1 contributes to AMPAR trafficking underlying mGluR-LTD. The presented experimental confirmation of candidate regulators in mGluR-LTD furthermore underlines the strength of this quantitative and high-resolution proteomics approach.

**Figure 6.**
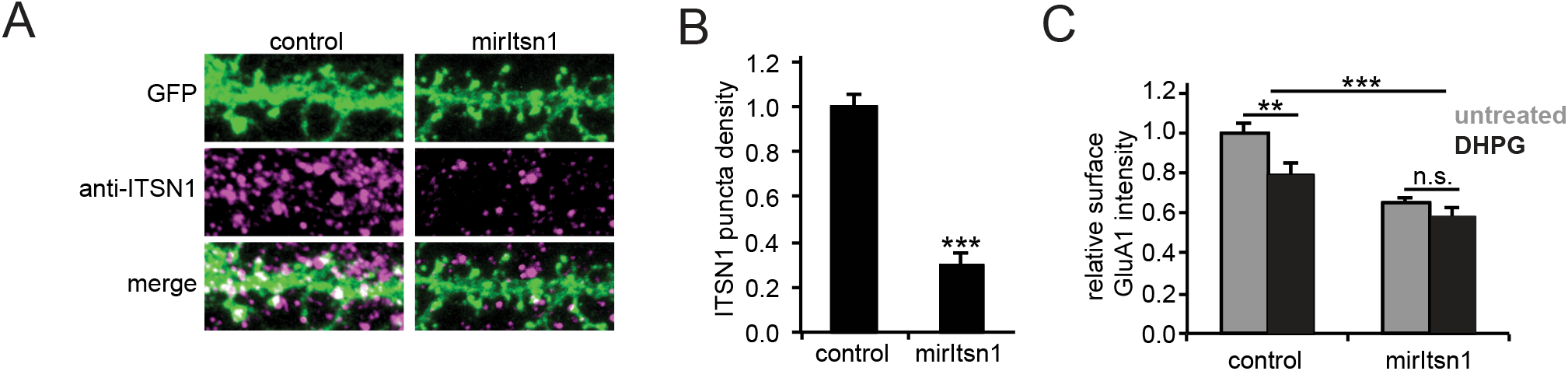
Itsn1 is essential for DHPG-induced AMPAR internalization. (A) Immunostaining of GluA1 of transfected neurons with a miRNA-based knockdown construct targeting both the long and short forms of Itsn1 (mirItsn1) or control transfection. (B). Immunostaining of endogenous Itsn1 confirms significant depletion of Itsn1 in mirItsn1-transfected neurons. (C) Surface expression of GluA1 was significantly reduced in Itsn1 knockdown neurons under basal conditions and GluA1 surface levels in response to DHPG are severely affected in Itsn1 knockdown neurons. Data are represented as mean±SEM ** p<0.01, *** p<0.001, n.s. not significant.

## Discussion

Here we used a combination of pulsed AHA and TMT labeling approaches to study protein synthesis during mGluR-LTD. This combination ensured for enrichment and MS-based relative quantification of labeled and thus newly synthesized proteins. This enrichment method has been shown to be applicable to different types of cultured cells, including primary cultures like neurons (Dieterich, Link et al. 2006, Eichelbaum, Winter et al. 2012, Kenney, Genheden et al. 2016, Schanzenbacher, Sambandan et al. 2016). A major advantage of the use of these pulsed labeling approaches in combination with mass spectrometry analysis is the possibility to identify and quantify subtle alterations in expression of proteins amidst the background of a steady state proteome, even of low abundant proteins. This approach led to the identification of 273 newly synthesized proteins upon mGluR activation, containing both known and novel proteins linked to mGluR-LTD. Here, we identified several proteins previously shown to be involved in mGluR-LTD, such as ERK1 (Thomas, G. M., Huganir 2004) and CamKII, and proteins implicated in endocytosis such as Cdc42 (Ramakers, G. J., Wolfer et al. 2012) and clathrin. Unfortunately, we did not identify some known mGluR-LTD markers including Arc/Arg3.1, Step or Ophn1 (Waung and Huber 2009; Luscher and Huber 2010), which might be caused by their expression abundance, the chosen time points of analysis, or the nature of the enrichment method. This could potentially be improved by higher affinity click chemistry enrichment techniques or fractionation before nanoLC-MS/MS analysis. For instance, CaMKIIa was identified as newly synthesized after DHPG stimulation in the proteomics data set but was not included for further analysis due to missing quantitative values. Further, we observed translation of several proteins that are themselves involved in protein synthesis, such as translation initiation and elongation factors, and multiple ribosomal proteins, but also proteins involved in proteasomal degradation, *e.g.* multiple proteasome subunits and enzymes involved in ubiquitination. These findings hint to the importance of regulating the fine balance between translation and degradation of proteins involved in the induction of mGluR-LTD.

Functional clustering of translated proteins upon induction of mGluR-LTD led to the identification of known LTD proteins, and highlighted several functionally relevant proteins with potential roles in mGluR-LTD related processes. Importantly, these functional families of proteins are not limited to the provided clusters, since these are solely based on annotated proteins with known functions or interactions. For instance, the splicing factors in cluster 1 (Figure 2), could potentially be supplemented with less-studied family members, including Srsf1, Srsf4, and Srsf7. This also holds true for, among others, proteasomal subunits and ribosomal proteins. These extended clusters and their corresponding heatmaps can be found in figure S5.

We complemented the protein translation dataset information on the phosphorylation events that potentially drive the signaling pathways that regulate mGluR-LTD. These events are usually very fast, to allow the cell to rapidly respond to external signals and precede protein translation. This makes phosphorylation likely the first step in the induction of mGluR-LTD, and it could therefore yield valuable information on the underlying molecular events, including receptor dynamics. Here, we confirmed the role of CaMKII in mGluR-LTD in both our sequence motif analysis and kinase inhibitor assay on GluA1 internalization. Multiple studies have shown the importance of kinase and phosphatase activity in the induction and maintenance of mGluR-LTD in general (Gallagher, Daly et al. 2004, Gladding, Fitzjohn et al. 2009, Hou, Klann 2004b), emphasizing the importance of phosphorylation in this dynamic process. Involvement of serine/threonine kinase activity has been studied intensively, and have been shown to be prominently involved in the activation of protein translation via the PI3K/Akt and subsequently Tsc and mTOR pathways, while tyrosine phosphatases and kinases are believed to be responsible for AMPAR tagging for internalization, and subsequent degradation (Bidinosti, Botta et al. 2016, O’Connor, Bariselli et al. 2014). Contrary to the intracellular signaling pathways that generally follow Ga_q/11_ protein stimulation, DHPG-LTD has repeatedly been shown to activate G-protein-independent signaling pathways (Fitzjohn, Palmer et al. 2001, Schnabel, Kilpatrick et al. 1999). In line with these findings, we find little phosphorylation, or translation, of proteins of the PLC, DAG, and PKC pathways leading to intracellular calcium release. Furthermore, the kinase inhibitor assay confirmed that inhibition of PKC activity does not influence DHPG-induced GluA1 internalization. This is in line with previous research, which also showed that DHPG induced mGluR-LTD is not dependent on PKC activity (Schnabel, Kilpatrick et al. 1999). Alternatively, we found significant regulation of CDK-type kinases. Cdk5 is a known regulator of mGluR5 activation, as it controls phosphorylation of the binding site of the adaptor protein Homer to the proline-rich C terminus of group I mGluRs. Via this mechanism, Cdk5 activation is negatively correlated with mGluR5 activation (Hu, Yang et al. 2012). Moreover, hippocampal slices treated with a Cdk4 inhibitor showed impaired DHPG-induced LTD (Li, C., Li et al. 2007), suggesting that at least two prominent members of the CDK family have functions in synaptic plasticity processes in hippocampal neurons. Our motif analysis indeed confirmed a role for CDK-type kinases in mGluR-LTD, as further demonstrated using the Cdk1, 2 and 5 specific inhibitor Roscovitine, which inhibited GluA1 internalization after DHPG stimulation, however, does not fully distinguish the individual roles of Cdk1, 2 and 5 in DHPG induced mGluR-LTD.

Next to kinase-specific signaling functions, clear regulation of cytoskeleton elements by phosphorylation was observed (Figure 3), suggesting cytoskeleton reorganization during mGluR-LTD. Most prominently, alterations in phosphorylation status of cytoskeletal regulators were observed, some of which were described before (Nakamura, Wood et al. 2011, Zhou, Hu et al. 2011), suggesting microtubule organization and actin reorganization. Next to cytoskeleton related processes, GO term analysis on molecular functions of the identified phosphorylated proteins yielded enrichment of regulatory activity of small GTPases. Several members of the Ras GTPase superfamily were found to be newly synthesized upon chemical mGluR-LTD induction, including members of the Rab and Ran subfamilies, Cdc42, and several of their interacting proteins. Recently, studies have been performed on some small GTPases in relation to several types of synaptic plasticity, shedding light on the possible importance of these types of molecules in mGluR-LTD as well. Overall, these and other data provide evidence for the possible importance of small GTPases in mGluR-LTD, and should be followed up further (Barnes, Wijetunge et al. 2015, Zheng, Jeyifous et al. 2015).

Itsn1 has not been subjected to extensive analysis in the context of synaptic plasticity before, although it was shown to influence trafficking of the AMPAR subunit GluA1 in *C. Elegans* (Glodowski, Chen et al. 2007). Here, we have shown a role for Itsn1 in GluA1 trafficking at dendritic spines in mammals as well. Interestingly, knockdown of Itsn1 reduced the expression of GluA1 at the dendritic spines, even in the absence of DHPG. Importantly, the induction of mGluR-LTD upon DHPG stimulation was omitted in the knockdown neurons, emphasizing the central role of Itsn1 in GluA1 internalization. Together this demonstrates a role for Itsn1 in GluA1 trafficking in general and more specific in mGluR-LTD induced GluA1 internalization. The exact mechanism by which Itsn1 influences receptor trafficking remains to be studied further. An interesting question is whether Itsn1 exerts its functionality mostly via posttranslational modifications such as the here identified phosphorylation sites, or via one of its interacting domains with other proteins. Previous research showed that its DH domain was critical for Cdc42 activation, and its SH3 domain for N-WASP interaction (Hussain, Jenna et al. 2001). Although one of our identified phosphorylation sites falls outside of these regions (S623 is located in the coiled coil part of the protein), the second one, S1134, falls within the N-WASP interacting SH3 domain.

In conclusion, we were able to construct a comprehensive map of signaling and translational events upon mGluR-LTD induction by integrating significant regulation of protein phosphorylation and translation. Over time, we could monitor activation of signaling pathways, as well as upregulation of protein signaling complexes involved in clathrin-mediated endocytosis. This multipronged analysis revealed several novel players in mGluR-LTD dependent AMPAR internalization, of which we could validate the involvement of Itsn1. We anticipate that our quantitative dataset on protein phosphorylation and translation during mGluR-LTD can be used as a rich resource for further analyses.

## Acknowledgements

A.F.M.A. is supported by the Netherlands Organization for Scientific Research (NWO) through a VIDI grant (723.012.102). This work was partly supported by *Proteins@Work*, a program of the Netherlands Proteomics Centre financed by the Netherlands Organisation for Scientific Research (NWO) as part of the National Roadmap Large-scale Research Facilities of the Netherlands (project number 184.032.201). H.D.M. is supported by the NWO through a VENI grant (863.13.020), by the European Research Council (ERC starting grant 71601) and received support from a FEBS Return-to-Europe fellowship and a NARSAD Young Investigator Award. C.C.H. is supported by the NWO (NWO-ALW-VICI 865.10.010), the Netherlands Organization for Health Research and Development (ZonMW-TOP 91213017 and 91215084) and the European Research Council (ERC) (ERC Consolidator grant 617050).

## Author contributions

C.A.G.H.G. and R.P. designed and conducted proteomics sample preparation, and performed the quantitative proteomics experiments and analyzed the data; H.M.G. and L.C. designed and performed functional kinase blocker and knockdown experiments and analyzed the data. A.F.M.A. supervised the quantitative proteomics experimental setup and the mass spectrometry data. The figures were designed and assembled by R.P., C.A.G.H.G., H.M.G., A.F.M.A., and C.C.H. The manuscript was written by R.P., C.A.G.H.G., H.M.G., and A.F.M.A. with input from C.C.H. A.F.M.A. and H.M.G. supervised the project and coordinated the study. Authors declare no competing financial interests.

## Materials and methods

### Ethics statement

All animal experiments were performed in compliance with the guidelines for the welfare of experimental animals issued by the Government of The Netherlands. All animal experiments were approved by the Animal Ethical Review Committee (DEC) of Utrecht University.

### Neuronal cultures

Hippocampal cultures were prepared from embryonic day 18 (E18) rat brains as described in (Esteves da Silva et al., 2015). Dissociated neurons were plated on poly-L-lysine (30 µg/ml) and laminin (2 µg/ml) at a density of 200,000 neurons per well. Cultures were grown in Neurobasal medium (NB) supplemented with B27, 0.5 mM glutamine, 12.5 µM glutamate, and penicillin / streptomycin at 37**°**C/5% CO_2_. Neurons were transfected at DIV10-14 with indicated constructs using Lipofectamine 2000 (Invitrogen) and experiments were performed 5 – 7 days later.

### Phosphopeptide analysis

#### DHPG stimulation and protein digestion

At DIV14-17 neurons were stimulated with 100 µM DHPG for 0, 5, 10, or 20 minutes. Neurons were washed three times with PBS and harvested directly in 8 M Urea lysis buffer supplemented with phosphatase inhibitor (PhosSTOP, Roche) and protease inhibitor (cOmplete mini EDTA-free, Roche). Neurons were lysed at 4°C with the Bioruptor Plus (Diagenode) by sonicating for 15 cycles of 30 sec. Protein content was determined with a Pierce BCA protein quantification assay (Thermo Fisher). Equal amounts of protein were heated at 95°C for 5 minutes and then reduced (4 mM DTT) for 20 minutes at 56°C and alkylated (8 mM IAA) for 25 minutes in the dark at room temperature. The proteins were the digested with Lys-C (1:75, Wako) for 4h at 37°C, after which the samples were diluted to a urea concentration of 2M and trypsin (1:50, Sigma Aldrich) was added overnight. The peptides were acidified to a total concentration of 1% Formic Acid (Merck). Samples were cleaned up using OASIS sample cleanup cartridges (Waters) and dried *in vacuo*.

#### Phosphorylated peptide enrichment

Phosphorylated peptides were enriched using Fe(III)-NTA cartridges (Agilent technologies) in an automated fashion using the AssayMAP Bravo Platform (Agilent technologies). The cartridges were primed with 0.1% TFA in ACN and equilibrated with loading buffer (80% ACN/0.1% TFA). Samples were suspended in loading buffer and loaded onto the cartridge. The peptides bound to the cartridges were washed with loading buffer and the phosphorylated peptides were eluted with 1% ammonia directly into 10% formic acid. Samples were dried *in vacuo* and stored at −80 °C until LC-MS/MS analysis.

#### Mass spectrometry and data-acquisition

The phosphorylated peptide enriched samples were analyzed with an UHPLC 1290 system (Agilent technologies) coupled to an Orbitrap Q Exactive Plus mass spectrometer (Thermo Scientific). Before separation peptides were first trapped (Dr Maisch Reprosil C18, 3 μm, 2 cm x 100 μm) and then separated on an analytical column (Agilent Poroshell EC-C18, 2.7 μm, 50 cm x 75 μm). Trapping was performed for 10 min in solvent A (0.1% FA) and the gradient was as follows; 4 - 8% solvent B (0.1% FA in acetonitrile) in 2 min, 8 - 24% in 71 min, 24 - 35% in 16 min, 35 - 60% in 7 min, 60 - 100% in 2 min and finally 100 % for 1 min. Flow was passively split to 300 nl/min. The mass spectrometer was operated in data-dependent mode. At a resolution of 35.000 *m/z* at 400 *m/z*, MS full scan spectra were acquired from *m/z* 375–1600 after accumulation to a target value of 3e^6^. Up to ten most intense precursor ions were selected for fragmentation. HCD fragmentation was performed at normalised collision energy of 25% after the accumulation to a target value of 5e^4^. MS/MS was acquired at a resolution of 17,500. Dynamic exclusion was enabled with an exclusion list of 500 and a duration of 18s.

#### Data analysis

RAW data files were processed with MaxQuant (v1.6.0.1(Cox, Mann 2008)) and MS2 spectra were searched with the Andromeda search engine against the TrEMBL protein database of Rattus Norvegicus (28,080 entries, downloaded 08/08/2017) spiked with common contaminants. Cysteine carbamidomethylation was set as a fixed modification and methionine oxidation, protein N-term acetylation, and phosphorylation of serine, threonine, and tyrosine were set as variable modifications. Trypsin was specified as enzyme and up to two miss cleavages were allowed. Filtering was done at 1% false discovery rate (FDR) at the protein and peptide level. Label-free quantification (LFQ) was performed, and “match between runs” was enabled. The data was further processed using Perseus 1.6.0.7 (Tyanova, Temu et al. 2016), WebLogo (Crooks, Hon et al. 2004, Schneider, Stephens 1990), MotifX (Park, Park et al. 2008, Pechstein, Bacetic et al. 2010, Chou, Schwartz 2011, Schwartz, Gygi 2005), and SynGO (Koopmans, van Nierop et al. 2019).

### pAHA & TMT labeling

#### Stimulation and lysis

DIV12 hippocampal neurons were incubated in NB media (Gibco life technologies) supplemented with B27, 0.5 µM glutamine and penicillin/streptomycin (supplemented NB) and either with 4 mM L-azidohomoalanine (L-AHA) (Bachem) or 4 mM L-Methionine (Sigma-Aldrich) and in parallel stimulated with 100 µM DHPG or vehicle for 5 minutes at 37**°**C/5% CO_2_. Neurons were then moved into freshly supplemented NB media with either 4 mM L-AHA or 4 mM L-methionine and incubated at 37**°**C/5% CO_2_ until the end of the experiment (15, 45, and 90 minutes after initial DHPG stimulation). Harvest followed three washes with PBS, directly into urea lysis buffer (Click-it Protein enrichment kit, Invitrogen C10416) supplemented with protease inhibitor (cOmplete mini EDTA-free, Roche).

#### Enrichment and digestion

Neurons were lysed at 4°C with the Bioruptor Plus (Diagenode) by sonicating for 10 cycles of 30 sec. Protein content was determined with a Pierce BCA protein quantification assay (Thermo Fisher). Newly synthesized proteins were then enriched using the Click-it protein enrichment kit for chemistry capture of azide modified proteins (Invitrogen C10416) following the manufactures protocol with small modifications. In short, protein lysate volume was adjusted to equal protein input for each sample per biological replicate and final volume was adjusted by adding Milli-Q. Lysates were added to washed resin and was incubated end to end rotating overnight. Resins were washed with Milli-Q and SDS buffer was added. Proteins were reduced (1M DTT) for 15 minutes at 70°C and alkylated (40 mM IAA) for 30 minutes in the dark at room temperature. The resins were then transferred to the supplied filter columns and extensively washed with subsequently SDS wash buffer, 8 M urea with 100 mM Tris pH 8, 20% ACN and 50 mM Ammonium Bicarbonate (AMBIC). Resins were transferred to a new tube and proteins were digested with 0.1 µg Lys-C for 2h at 37°C and 0.5 µg trypsin (Promega) overnight at 37°C. Samples were centrifuged and the supernatant was taken. Sample cleanup was performed using the OASIS sample cleanup cartridges (Waters). Samples were dried *in vacuo* and stored at −80 °C.

#### TMT labeling

TMT labeling was performed according to the manufacturer’s instructions using the TMT10plex Isobaric Label Reagent Set (Thermo Scientific). In brief, samples were reconstituted in 100 µl 87.5% HEPES buffer pH 8.5 / 12.5% ACN, and TMT reagents in 41 µl anhydrous ACN. Full contents of the reagents were added to the samples and incubated at room temperature for one hour, after which the reaction was quenched with 5% hydroxylamine. TMT reagents 126 – 130C were used to label the nine experimental conditions, and these were mixed in a 1:1 ratio. The TMT 131 label served as a reference ‘pool’ between biological replicates and consisted of an equal mix of all nine experimental conditions of biological replicate 1, and was mixed in equal ratios with the other TMT labels. Labeled mixtures were cleaned using the OASIS sample cleanup cartridges (Waters), dried *in vacuo* and stored at −80 °C until further processing.

#### Mass spectrometry and data-acquisition

Fractions were reconstituted in 10% FA and analyzed in two technical replicates with a UHPLC 1290 system (Agilent technologies) coupled to an Orbitrap Q Exactive X mass spectrometer (Thermo Scientific). Peptides were trapped on an in house made trap column (Dr Maisch Reprosil C18 column, 3 µm, 2 cm x 100 µm) and separated on an analytical column (Agilent Poroshell EC-C18, 2.7 µm, 50 cm x 75 µm). Trapping was performed for 5 min in solvent A (0.1% FA) and separation was performed using a 85 min linear gradient from 15% to 45% solvent B. Flow was passively split to 300 nl/min. The mass spectrometer was operated in data-dependent mode. At a resolution of 60,000 at 200 m/z, MS full scan spectra were acquired from 375 – 1600 m/z after accumulation to a target value of 3e^6^. Up to 15 most intense precursor ions were selected for HCD fragmentation at a normalized collision energy of 32% after accumulation to a target value of 5e^4^. MS/MS was acquired at a resolution of 60,000, with a fixed first mass of 120 m/z. Dynamic exclusion was enabled with a duration of 12s.

#### Data analysis

RAW data files were processed using Thermo Proteome Discoverer (version 2.2.0.338) and Mascot search engine (v2.6.1), allowing for variable methionine oxidation, protein N-terminal acetylation, and methionine replacement by AHA. Carbamidomethylation of cysteines was set as a fixed modification. The protein database consisted of the TrEMBL protein database of Rattus Norvegicus (28,080 entries, downloaded 08/08/2017) spiked with common contaminants. Enzyme specificity was set for trypsin, with a maximum of two allowed missed cleavages. The precursor mass tolerance was 50 ppm, and fragment mass tolerance was set to 0.05 Da. TMT 10plex was set as quantification method, and only unique peptides were used for quantification. Normalization mode was disabled, and reporter abundances were based on signal to noise values in all cases.

### Data availability

The mass spectrometry proteomics data have been deposited to the ProteomeXchange Consortium via the PRIDE partner repository with the dataset identifier PXD014043.

### DNA constructs

The Itsn1 miRNA knockdown construct was generated by annealing oligos containing the 21-nucleotide targeting sequences described in (Thomas, S., Ritter, Verbich, Sanson, Bourbonniere, McKinney, and McPherson 2009b) and ligating in the miRNA expression plasmids pSM155-GFP (provided by G. Du; University of Texas, Houston, TX) (Du et al., 2006) digested with BsmBI.

### Kinase inhibitor assay

For the kinase inhibitor assays, neurons were pre-incubated with KN-93 (10 µM), staurosporine (1 µM), or roscovitine (20 µM) for 30 minutes before induction of mGluR-LTD. Neurons were stimulated with DHPG (100 µM) for 5 minutes, returned to original medium and fixed 30 minutes later. Blockers were present during the entire experiment.

### Immunofluorescence and confocal microscopy

To induce mGluR-LTD, hippocampal neurons were stimulated with DHPG for 5 minutes and then returned to the original culture medium. After 30 minutes, neurons were fixed with 4% paraformaldehyde / 4% sucrose in phosphate-buffered saline (PBS) for 8 – 10 minutes. Fixed neurons were blocked with 10% normal goat serum in PBS for 30 – 60 minutes at room temperature, stained with rabbit anti-GluA1 (1:100; Calbiochem), or anti-ITSN1 and labeled with fluorescent goat anti-rabbit secondary antibodies. Confocal images were taken with a Zeiss LSM 710 with 63x 1.40 oil objective. Images consist of a z-stack of 7-9 planes at 0.39 µm interval, and maximum intensity projections were generated for analysis and display. GluA1 cluster intensity was measured using the ParticleAnalyzer function in ImageJ and analyzed per region of interest.

---

**Figure S1.**
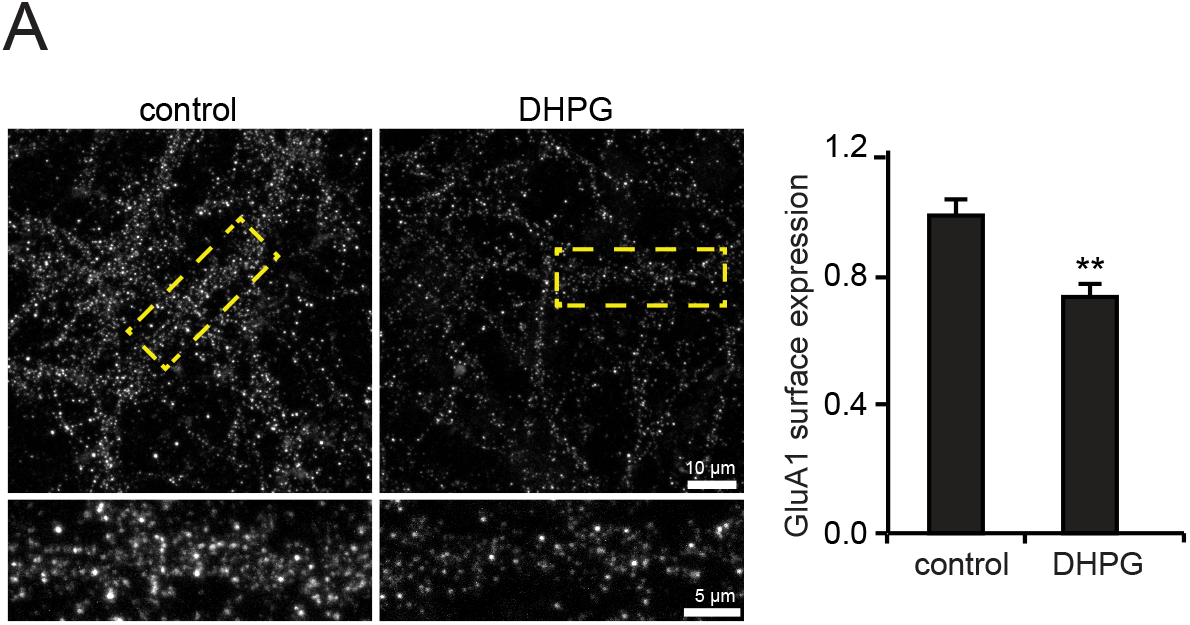
DHPG induces mGluR-LTD. (A) DHPG stimulation induces significant GluA1 internalization. Data are represented as mean±SEM. ** p<0.01.

**Figure S2.**
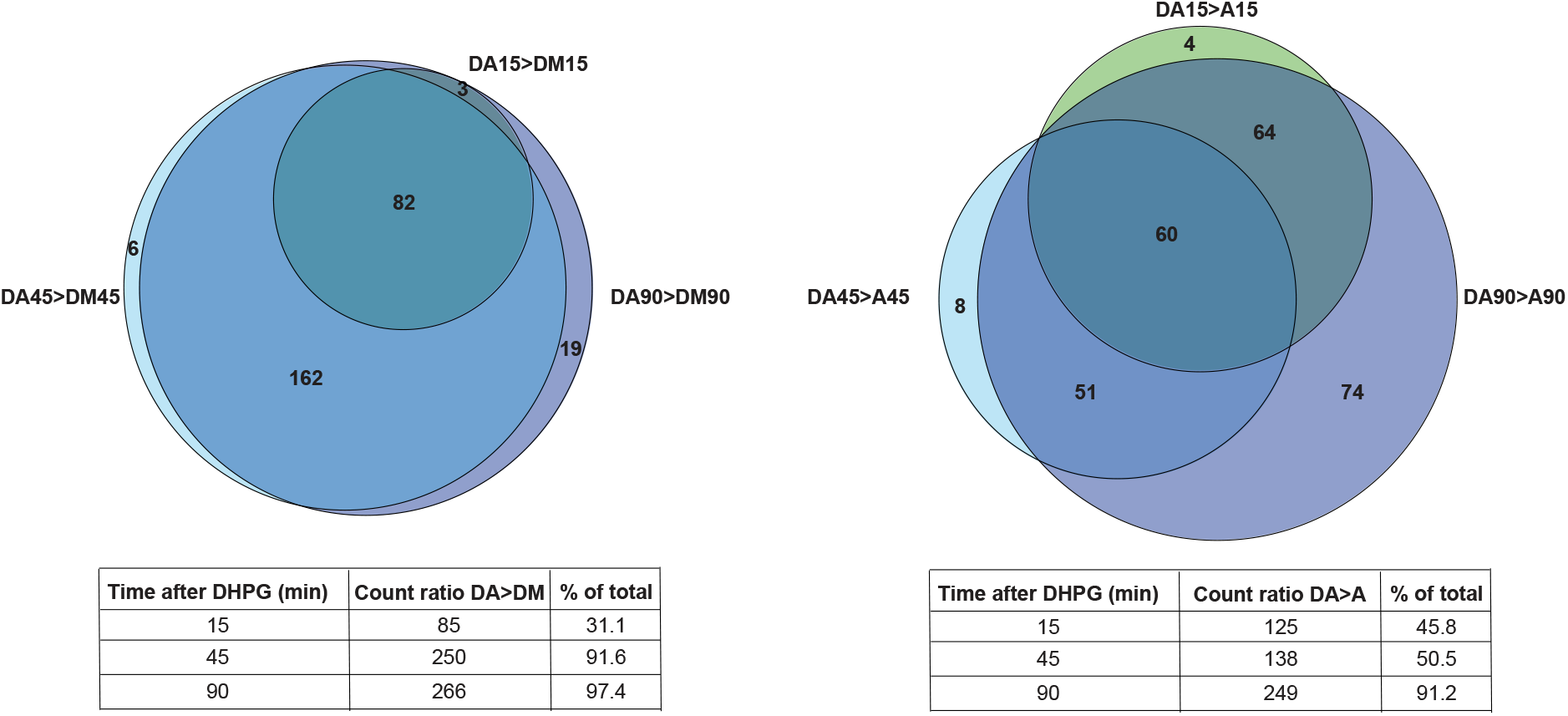
Evaluation of the AHA and TMT dataset. Venn diagrams showing the overlap of proteins identified in the different experimental conditions. As expected, the number of translated proteins increase over time, and the vast majority of proteins identified are stably identified among all studied time points. DA – DHPG and AHA, DM – DHPG and methionine, A – AHA only.

**Figure S3.**
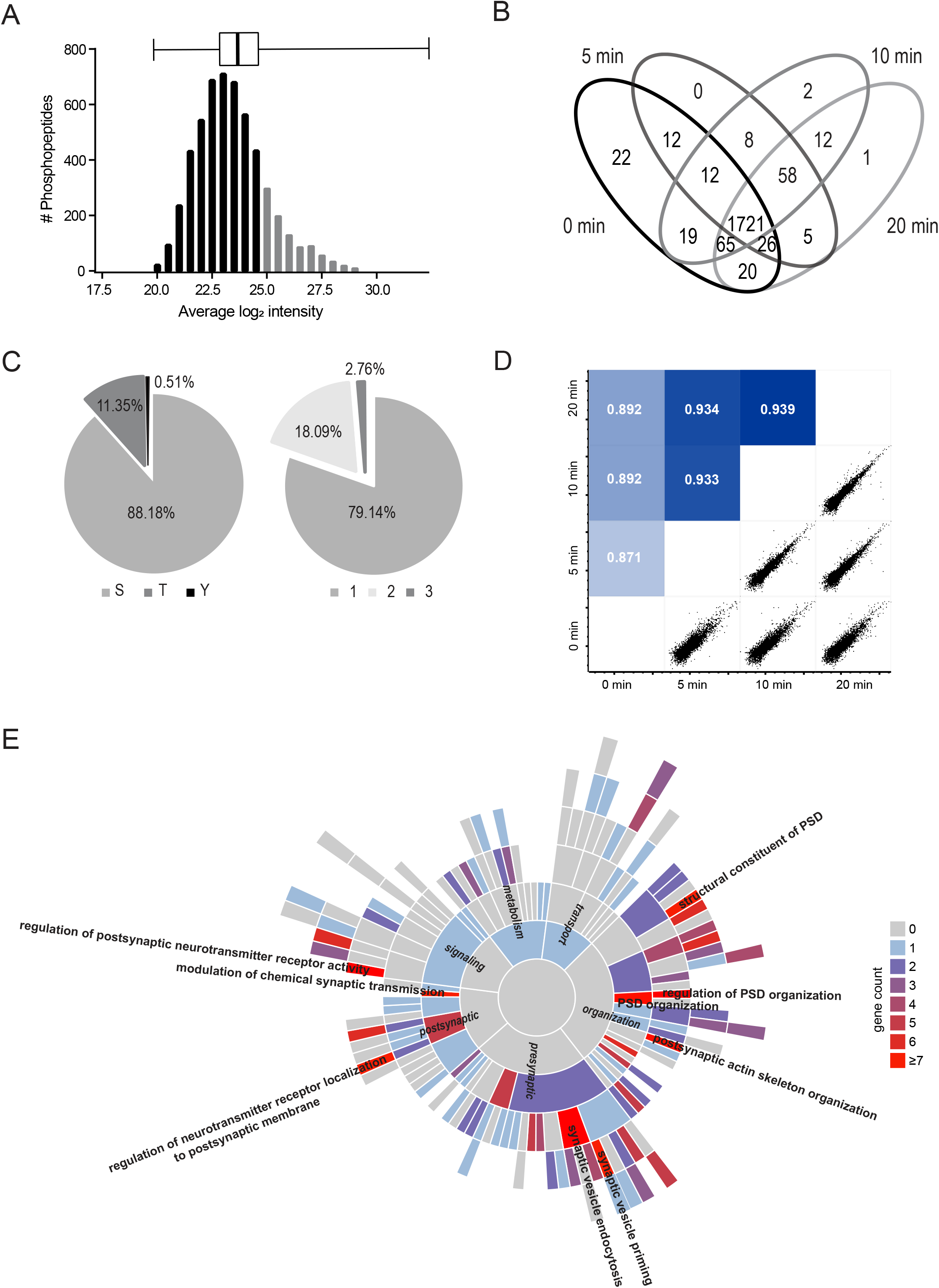
Evaluation of the phosphoproteomics dataset. (A) Distribution of phosphopeptide abundance, displaying normal distribution. Light grey bars indicate the top 25% most abundant phosphopeptides. (B) Venn diagram of the overlap between proteins identified in the 5, 10, and 20 minutes LTD experiment, as well as the control condition. Biological replicates were combined. (C) Percentages of enriched serine, threonine, and tyrosine phosphosites, and the distribution of singly, doubly and triply phosphorylated peptides. (D) Heatmap of Pearson correlations and correlation plots for the different biological replicates in the DHPG stimulated and control samples, showing high quantitative reproducibility between all measurements. (E) SynGO enrichment analysis of biological process. Processes with highest gene counts are labeled.

**Figure S4.**
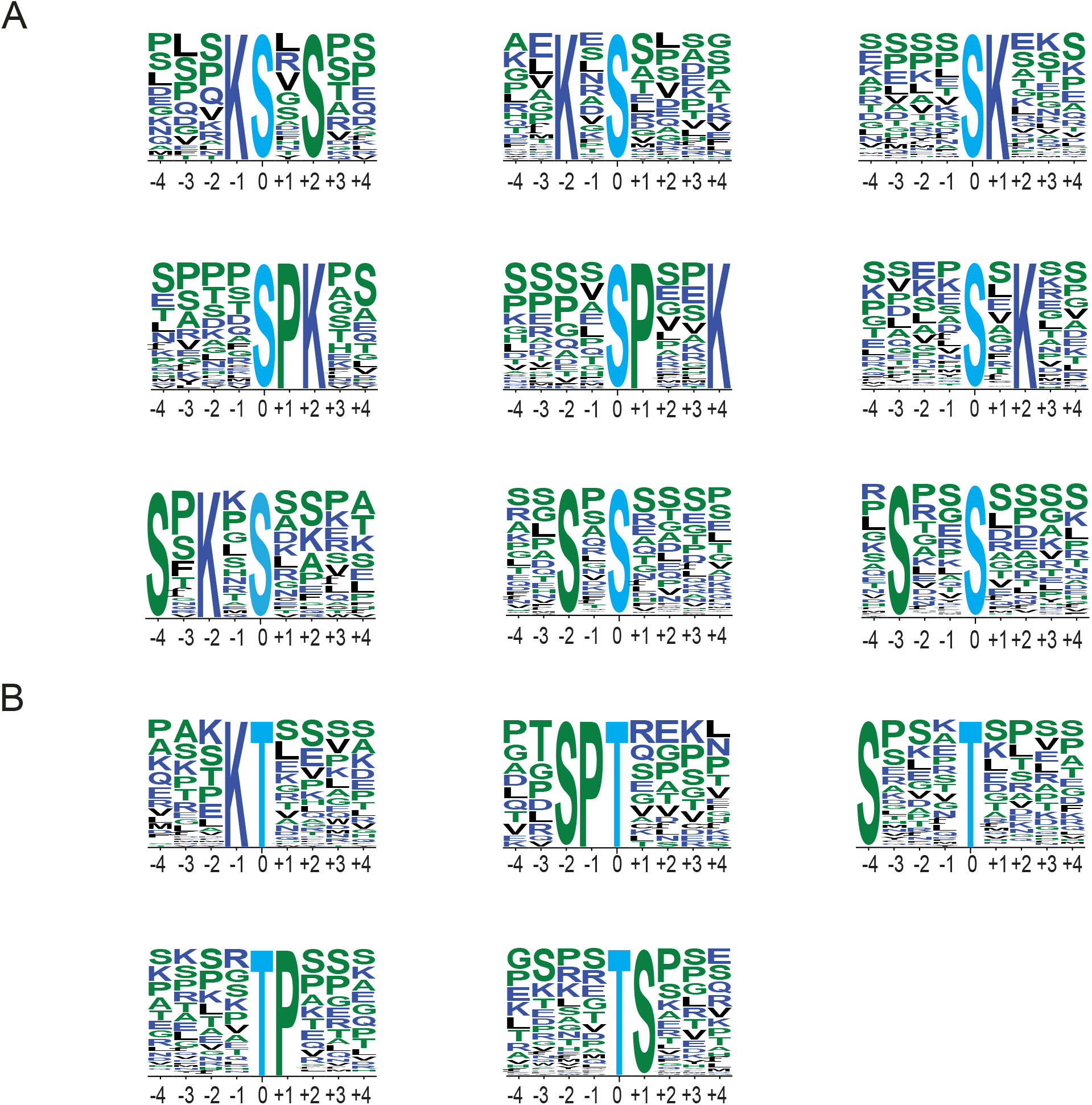
Phosphorylation sequence motifs. (A) Significantly enriched serine-directed phosphorylation motifs as generated with MotifX. (B) Significantly enriched threonine-directed phosphorylation motifs.

**Figure S5.**
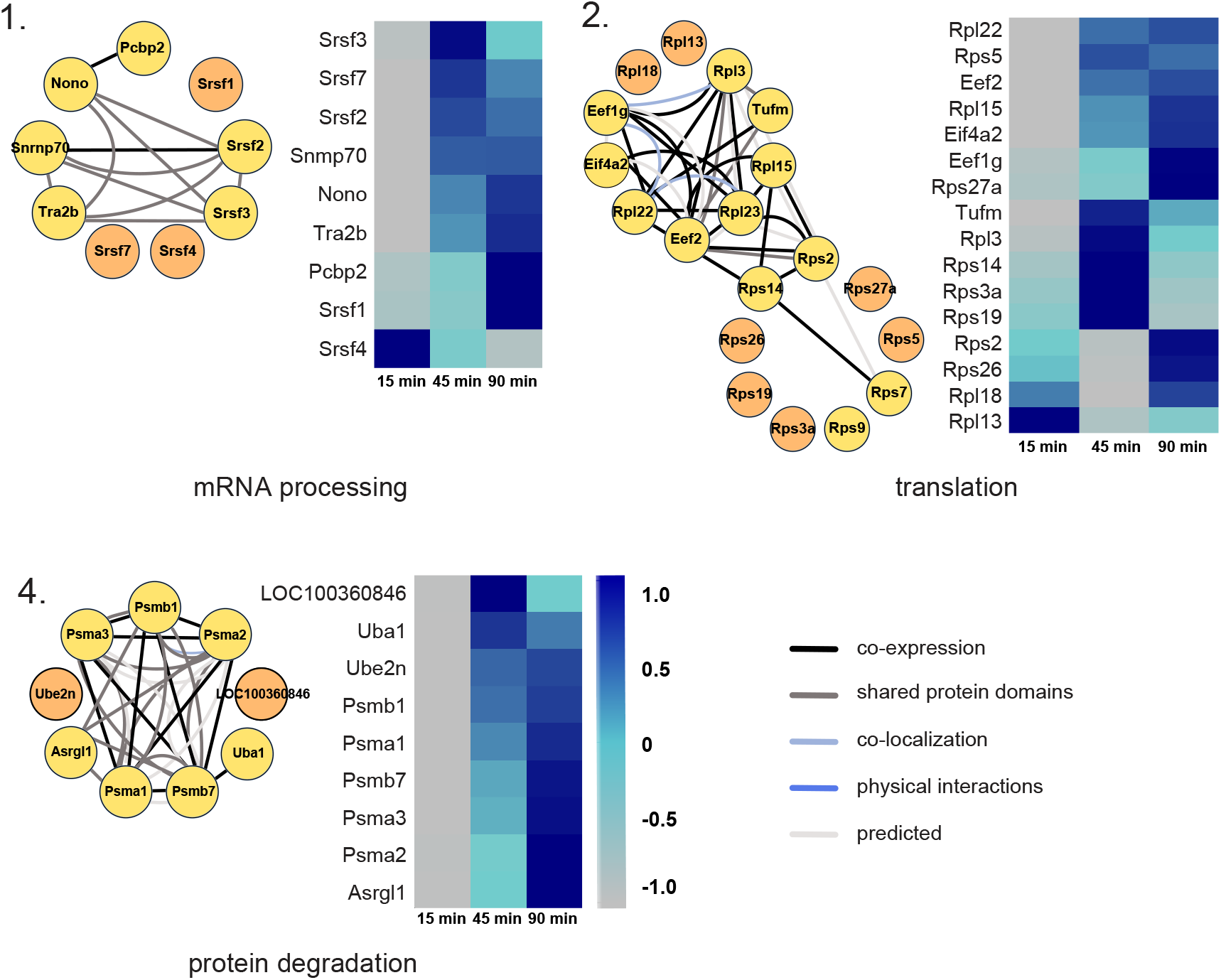
Supplemented interaction-based protein clusters of newly translated proteins. Protein clusters of enriched GO terms are displayed with their known interaction profiles (yellow), supplemented with less studied proteins from the translation dataset with similar function (orange). Heatmaps represent the z-score normalized ratio DHPG AHA / pool over the three measured time points.

**Table S1.** Overview of *de novo* synthesized proteins upon DHPG stimulation.

**Table S2.** Significantly regulated phosphorylation sites upon DHPG stimulation.

